# Minimally invasive delivery of peptides to the spinal cord for behavior modulation

**DOI:** 10.1101/2022.05.20.492752

**Authors:** Zhenghong Gao, Eric T. David, Tiffany W. Leong, Xiaoqing Li, Qi Cai, Juliet Mwirigi, Monica Giannotta, Elisabetta Dejana, John Wiggins, Sharada Krishnagiri, Robert M. Bachoo, Theodore J. Price, Zhengpeng Qin

## Abstract

The blood-spinal cord barrier (BSCB) tightly regulates molecular transport from the blood to the spinal cord. Herein, we present a novel approach for transient modulation of BSCB permeability and localized delivery of peptides into the spinal cord for behavior modulation with high spatial resolution. This approach utilizes optical stimulation of vasculature-targeted nanoparticles and allows delivery of BSCB-nonpermeable molecules into the spinal cord without significant glial activation or impact on animal locomotor behavior. We demonstrate minimally invasive light delivery into the spinal cord using an optical fiber and BSCB permeability modulation in the lumbar region. Our method of BSCB modulation allows delivery of bombesin, a centrally-acting and itch-inducing peptide, into the spinal cord and induces a rapid and transient increase in itching behaviors in mice. This minimally invasive approach enables behavior modulation without genetic modifications and is promising for delivering a wide range of biologics into the spinal cord for behavior modulation and potentially therapy.

**Significance Statement:** Spinal cord diseases and disorders are common and cause significant disability, including chronic pain, paralysis, cognitive impairment, and mortality. The blood-spinal cord barrier is a considerable challenge for delivery by systemic therapeutic administration. We developed an optical approach for effectively and safely delivering molecules to the spinal cord to overcome this barrier. The fiberoptic method is minimally invasive and overcomes challenges that previous technologies face, including the complicated bone structure and standing waves that complicate BSCB opening using ultrasound. Optical stimulation offers unprecedented spatial resolution for the precise delivery in intricate spinal cord structures. Significantly, our approach modulates animal behavior (i.e., itch) without genetic modifications and demonstrates the potential for delivery of biologics such as peptides into the spinal cord.

## Introduction

Neurological diseases and disorders affecting the spinal cord, such as spinal cord injury (SCI), amyotrophic lateral sclerosis (ALS), and chronic pain, have a significant societal impact (1). Researchers have investigated a wide range of potential therapeutics to treat spinal cord diseases and disorders (2). However, clinical trials for neurological diseases and conditions have limited therapeutic success partly due to insufficient delivery (3). One common challenge is the difficulty in delivering biologics or large molecules to the spinal cord due to the blood-spinal cord barrier (BSCB) (4, 5). Like the blood-brain barrier in the brain, the BSCB prevents 98% of large molecules from entering the spinal cord and only allows entry of a small number of molecules (glucose, oxygen, amino acids, and certain small, lipophilic compounds) (5, 6).

The BSCB is a unique structure in the spinal cord. It includes endothelial cells, tight junctions, basal membrane, and astrocyte end-feet that collectively constitute a functional neurovascular unit and highly regulated barrier (4, 7). The BSCB safeguards the immune-privileged CNS from toxins, pathogens, and other potentially harmful substances in the blood, thus playing a pivotal role in maintaining the optimal environment for neuronal signaling. At the same time, the BSCB is also a significant roadblock to the effective treatment of diseases and disorders in the spinal cord (5). Therefore, there is a critical need for novel approaches to overcome the BSCB for delivery to local spinal cord regions for therapeutic and neuromodulation applications (6).

There are several methods to overcome the BSCB (8). Transient opening of the BSCB mediated by focused ultrasound (FUS) following intravenous administration of gas-filled microbubbles (MBs) is currently an area of intense interest in improving spinal cord drug delivery (6, 9-12), as encouraged by promising results in the brain (6, 13). However, this technology may face many challenges in forming a tight ultrasound focus within the spinal canal, mainly due to the standing waves created by the reflective walls of the spinal canal, acoustic-thermal deposition in the bone (14), and complicated spine geometry (7). Intrathecal delivery has been used to administer therapeutics to the spinal cord (15). However, the CSF flow dynamics and administered molecule affect the intrathecal drug distribution. It remains a significant challenge to achieve localized spinal cord delivery.

Light can be delivered to specific locations in the spinal cord with high spatiotemporal resolution. Recent studies have demonstrated implanted optoelectronic systems in the epidural space for optogenetic neuromodulation in the spinal cord (16). However, the use of genetic transfection could impede its further clinical translation. Here, we present a new method that enables BSCB modulation (Opto-BSCB) and peptide delivery to the spinal cord for behavior modulation in mice. This method relies on the optical stimulation of vascular-targeted gold nanoparticles by short-pulsed laser light (17). For the first time, we demonstrate the local delivery of biologics (i.e., a peptide) to the spinal cord and robust itch behavior modulation in a rodent model with a minimally invasive optic fiber and without significant motor behavioral issues. Compared with other approaches such as focused ultrasound, Opto-BSCB allows direct light delivery to the spinal cord to modulate BSCB permeability via a fiberoptic device, bypassing the vertebrae with minimal invasiveness. Opto-BSCB opens many opportunities for improving the local concentration of a wide range of molecules, biologics, and therapeutic agents in the spinal cord.

## Results

### Optical BSCB modulation with a free-space laser beam

First, we tested the BSCB opening with laser stimulation of vasculature-targeted nanoparticles in vivo. We prepared vasculature-targeted nanoparticles by coating the nanoparticle surface with anti-JAM-A antibodies (BV11), followed by polyethelyne glycol (PEG) backfilling (18). We prepared control nanoparticles with PEG coating and without antibody functionalization. Transmission electron microscopy (TEM) shows the uniform size distribution centered at 48 nm (48 ± 9.2nm) (Fig. S1A, and insert, see Supplementary Information Appendix for further supplementary figures), consistent with the size measured by Dynamic Light Scattering (DLS) (Fig. S1B). The hydrodynamic radius increased to 70 nm after BV11 antibody functionalization. The absorbance spectrum measurement showed a strong absorption peak at 528 nm for AuNP-PEG (Fig. S1C), and it shifted to 532 nm after antibody conjugation for AuNP-BV11 (Fig. S1C). To detect whether the AuNP-BV11 can target the spinal cord, we then tested the biodistribution in mice after the intravenous (IV) injection of AuNP-PEG or AuNP-BV11 (Fig. S2) at 1 hour and two weeks post-injection. We found that AuNP-BV11 has significantly higher accumulation in the spinal cord when compared to control AuNP-PEG (6.5 times at 1 hour). We then intravenously injected AuNP-BV11 nanoparticles and stimulated the spinal cord using a laser beam through the skin (Fig. 1, 6 mm laser beam size, 40 pulses). Examination of Evans Blue extravasation suggests successful BSCB opening in the laser-irradiated area, while there was no blue dye leakage in the spinal cord regions without laser stimulation (Fig. 1C). Fluorescent imaging of the cross-section of spinal tissue further showed highly coordinated Evans Blue fluorescence distribution within the area of light stimulation but not in the region without light, confirming increased BSCB permeability (Fig. 1C). The Evans Blue fluorescence was about 20-fold higher for the area with laser applied than the region without light (p<0.0001, Fig. 1D).

**Figure 1.**
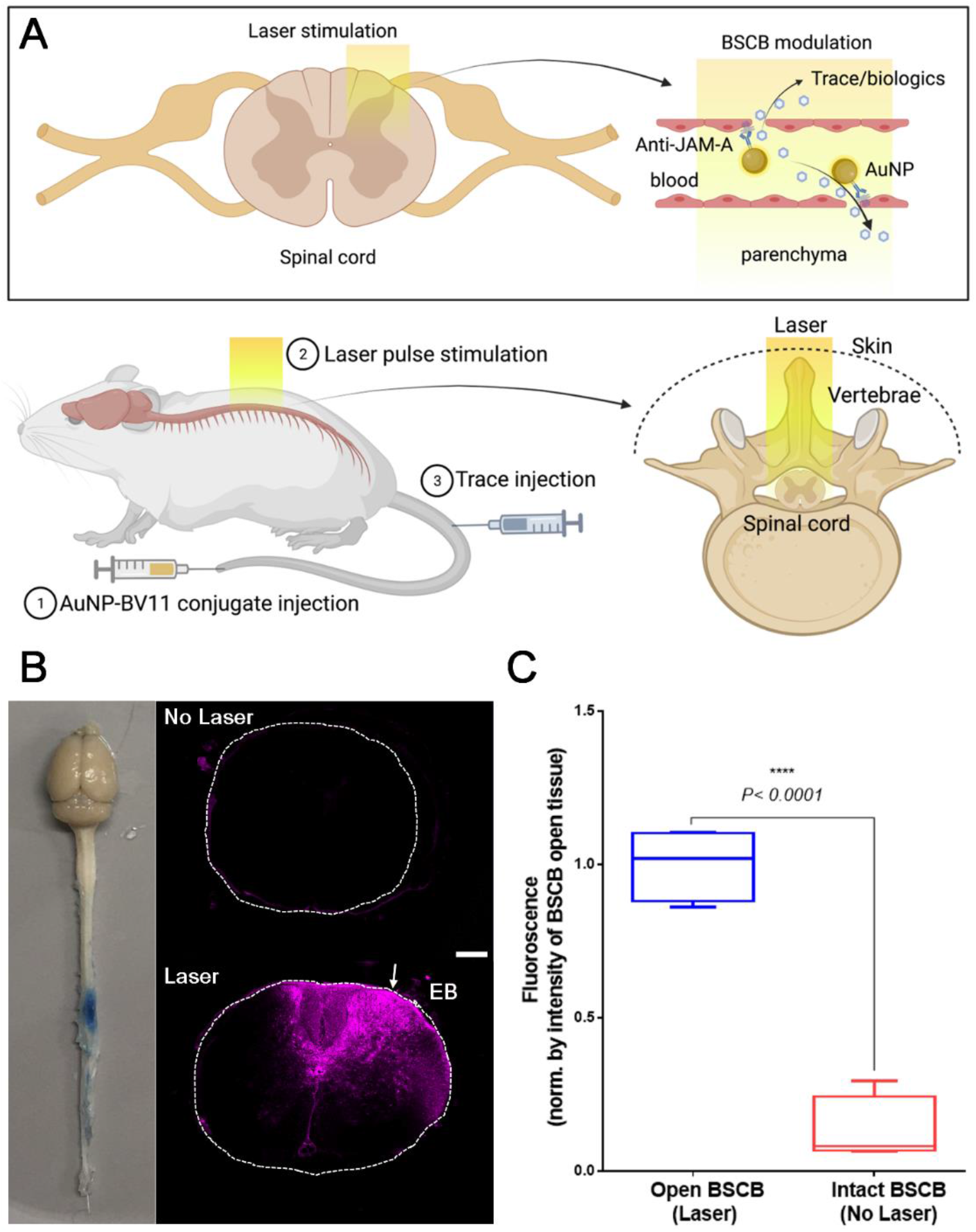
Opto-BSCB modulation with the bulk laser beam (beam size of 6 mm in diameter). (A) Schematic representation of the BSCB modulation via vasculature-targeted plasmonic AuNPs under laser pulse stimulation. (B) Whitefield and fluorescent images of the mouse CNS shows the Evans blue (EB) dye penetration in spinal tissue regions with (blue color) and without (natural color) laser stimulation. Laser conditions: 40 pulses at 5 Hz frequency and 35 mJ/cm^2^ laser power. Scale bar: 200 µm. (C) Quantitative comparison of the fluorescence in the spinal cord with open vs. intact BSCB shows a 20-fold higher fluorescence in the open BSCB tissue as compared with intact BSCB tissue (fluorescence is indicated as mean ± SD, normalized by the mean intensity at open BSCB tissue *N*=4, ****p<0.0001).

To demonstrate the versatility of our method of BSCB modulation, we directly pointed the laser beam at the skin covering the spine. We scanned half of the spinal region by moving the laser beam along the midline of the spine from the thoracic to the lumbar areas. (6 mm in diameter, 35 mJ/cm^2^, 10 pulses, frequency 5 Hz) We then carefully shaved the mouse skin to reduce light scattering, cleaned it with ethanol, and wetted it with water (Fig. S4). After nanoparticle administration and laser irradiation (6 mm in diameter, 35 mJ/cm^2^, 10 pulses), we observed Evans Blue leakage into the spinal cord in the laser beam-scanned region (Fig. S3). We noted no BSCB opening in the animals injected with AuNP-PEG under the same conditions (Fig. S4A-B) and no significant changes in the blood vessel morphology (Fig. S4C-D).

### BSCB modulation with fiberoptic light delivery

Next, we attempted fiberoptic light delivery using a small diameter (50-200 µm) optical fiber into the spinal cord. The optical fiber can be inserted into the spinal cord with a procedure similar to a clinically used lumbar puncture. To test whether the optical fiber delivers sufficient laser energy for BSCB modulation, we first performed a lumbar laminectomy by removing the tissue and vertebrae to expose the spinal cord. An optical fiber was then directly pointed at the surface of the tissue to deliver light (Fig. S5A). Upon AuNP-BV11 nanoparticle administration and laser stimulation (35 mJ/cm^2^, 5 pulses), we observed the Evans Blue dye leakage, as indicated by a small blue spot in the area with light stimulation (Fig. S5B-C). This observation suggests the highly localized BSCB modulation.

After validating the feasibility of fiberoptic light delivery, we used an intrathecal cannula to guide the optical fiber delivery with minimal invasiveness (Fig. 2). To study the BSCB opening size and duration, we IV injected fluorescent tracers including 600 Da EZ-link biotin and 70 kDa FITC-Dextran (Fig. 2A). EZ-link biotin and FITC-dextran could cross the BSCB to reach spinal tissue at 1 hour post-laser stimulation (Fig. 2B), which decreases significantly after 24 hours (Fig. 2C and 2E). This result demonstrates a reversible BSCB opening within a 24-hour window, consistent with findings in the brain (17). We then investigated the effect of laser pulse number on BSCB opening to unveil the potential for tunable delivery (Fig. 2D). By increasing pulse numbers from 1 to 20, the area with both EZ-link biotin and FITC-dextran leakage increased, saturating at 20 pulses. Under all conditions, the area covered by EZ-link biotin was greater than that of FITC-Dextran (Fig. 2F). The depth of dye leakage (from the surface of the spinal cord) increases when increasing the number of laser pulses from 1 to 10 (Fig. 2G). We observed significantly increased fluorescence intensity for FITC-Dextran and only slightly increased fluorescence intensity for EZ-link biotin when increasing the number of pulses from 1 to 40 (Fig. 2H). These results suggested that the laser pulse number can modulate the area of the BSCB opening. We further investigated the impact of BSCB opening on cell morphologies and cell number by immunohistochemistry (IHC) staining (Fig. 2I and J). We assessed main cell types in the spinal cord, including neurons (NeuN), astrocytes (GFAP), and microglial cells (Iba1). We observed no significant changes in NeuN, GFAP, or Iba1 staining at the site with laser light compared to regions with no light. Therefore, this approach does not disrupt neuronal integrity or cause significant astrogliosis or microglial reactivity changes.

**Figure 2.**
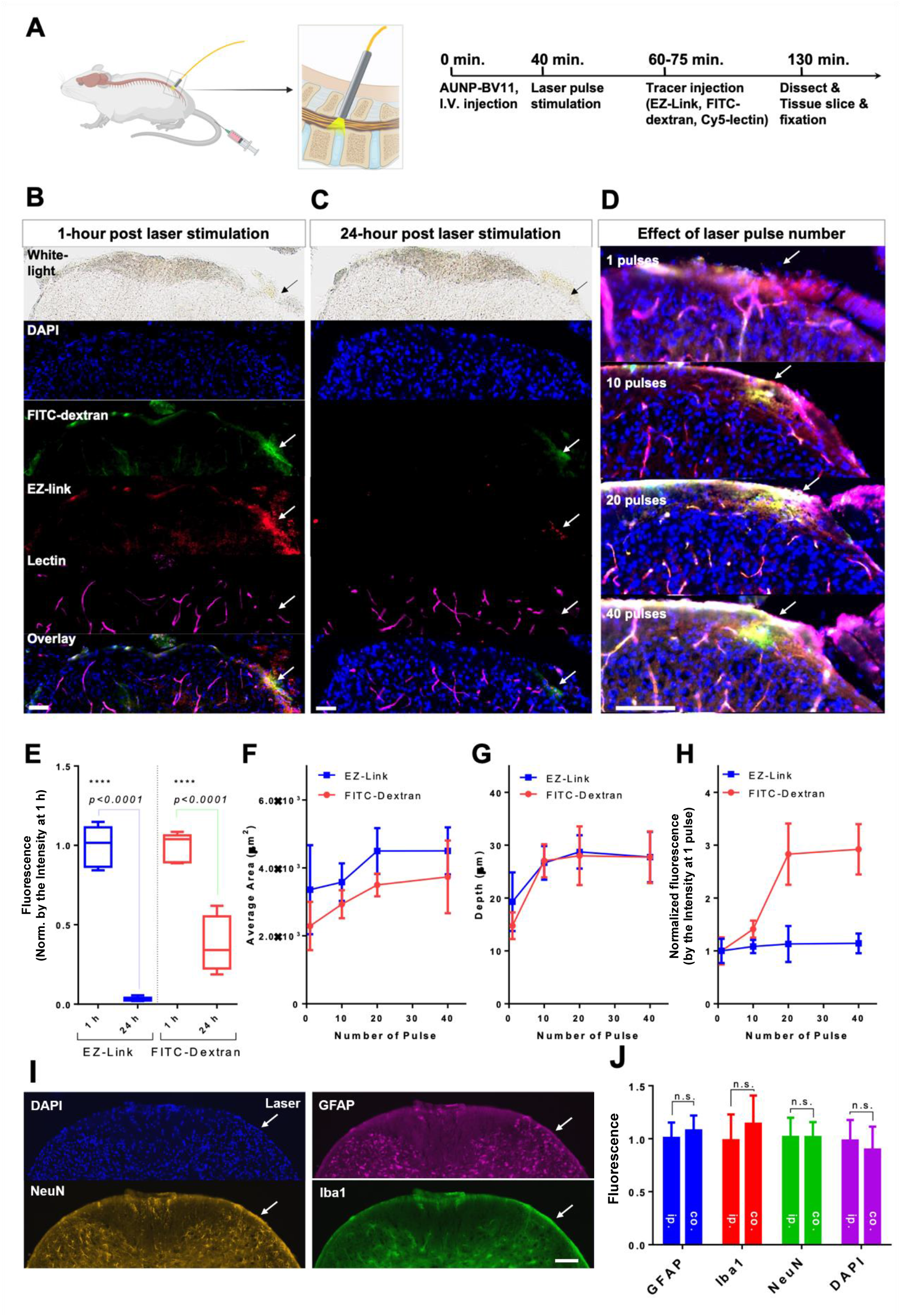
Minimally invasive Opto-BSCB modulation through fiberoptic light delivery. (A) Schematic of the fiberoptic light delivery for Opto-BSCB modulation and stepwise experimental protocol. (B-C) Permeability change of the BSCB post-laser stimulation at 1 hour and 24 hours. White light, DAPI, FITC-dextran 70kDa, EZ-link biotin (600 Da), lectin-Cy5, and overlay images were labeled for 1 and 24 hours. The white arrow represents the side of laser application. Scale bar: 50 µm. (D) Effect of laser pulses on BSCB permeability. IHC staining of the spinal cord tissue is performed by DAPI (nuclei, blue), EZ-link biotin is tagged by 2^nd^ antibodies labeled by Cy3 (red), and the blood vessels are identified by lectin-Cy5 (purple). Images from each color channel are overlayed. The white arrow represents the side of laser application. Scale bar: 50 µm. *N*=3 per group. (E) Quantitative comparison of the fluorescence of EZ-link biotin and FITC dextran administrated 1 hour and 24 hours after BSCB opening (fluorescence is indicated as mean ± SD, normalized by the mean intensity of 1 hour for both EZ-link and FITC-Dextran, *N*=3, p<0.0001 for both tracers, T-test). (F-H) Quantitative comparison of effect of pulse number on BSCB modulation: (F) The average fluorescent areas of EZ-link and FITC-dextran, (G) the depth, and (H) the intensity of fluorescence plotted as a function of pulse number increasing from 1, 10, 20, to 40 (intensity is indicated as mean ± SD, normalized by the mean at 1 pulse, *N*=3). Laser intensity: 35 mJ/cm^2^ at the fiber tip. (I) IHC staining of the spinal cord tissue by DAPI (nuclei), NeuN (neurons), GFAP (astrocytes), Iba1 (microglia). The white arrow represents the side of laser application. Scale bar: 100 µm. (J) Quantitative comparison of the fluorescence in IHC-stained tissues. Fluorescence is indicated as mean ± SD, normalized by the mean intensity at BSCB open side, *N*=4, NS: no significance, T-test. Ip: ipsilateral (intact BSCB, with no laser); co: contralateral (open BSCB, with laser).

### Minimally invasive and localized peptide delivery to the spinal cord

Lastly, we investigated the localized delivery of biologics to the spinal cord. We used a centrally-acting itch-inducing peptide (22, 23), bombesin (MW: 1.62 kDa), to test this idea. After systemic administration following the BSCB modulation (Fig. 3A), we observed a rapid and transient itching behavior in the mice with BSCB opening (Fig. 3B) 5-10 minutes post-injection of bombesin. In contrast, we observed no itching behavior above baseline in the mice with intact BSCB (Fig. 3B, and video S1). The increased itch score in mice with opened BSCB treated with bombesin lasted up to 10 minutes, consistent with the peptide clearance profile (19, 20). These results demonstrate the delivery of a small peptide to the spinal cord to introduce behavior-specific changes in a rodent model. We further investigated whether Opto-BSCB impacts other aspects of animal behavior, including motor function and paw temperature of the mice (Fig. 3C and D). We found no significant difference in the motor function assessed by the rotarod testing between open and intact BSCB mice (Fig. 3C). We assessed paw temperature using FLIR imaging, a non-invasive method used as a proxy for altered blood flow associated with inflammation. While we did not observe a significant change in paw temperature over time, both groups’ temperature drop suggested no inflammation induced by treatment (Fig. 3D). We attribute the decreased paw temperature over time to habituation. Together, these results indicate that the Opto-BSCB method would not likely alter animal motor coordination or lead to peripheral inflammation.

**Figure 3.**
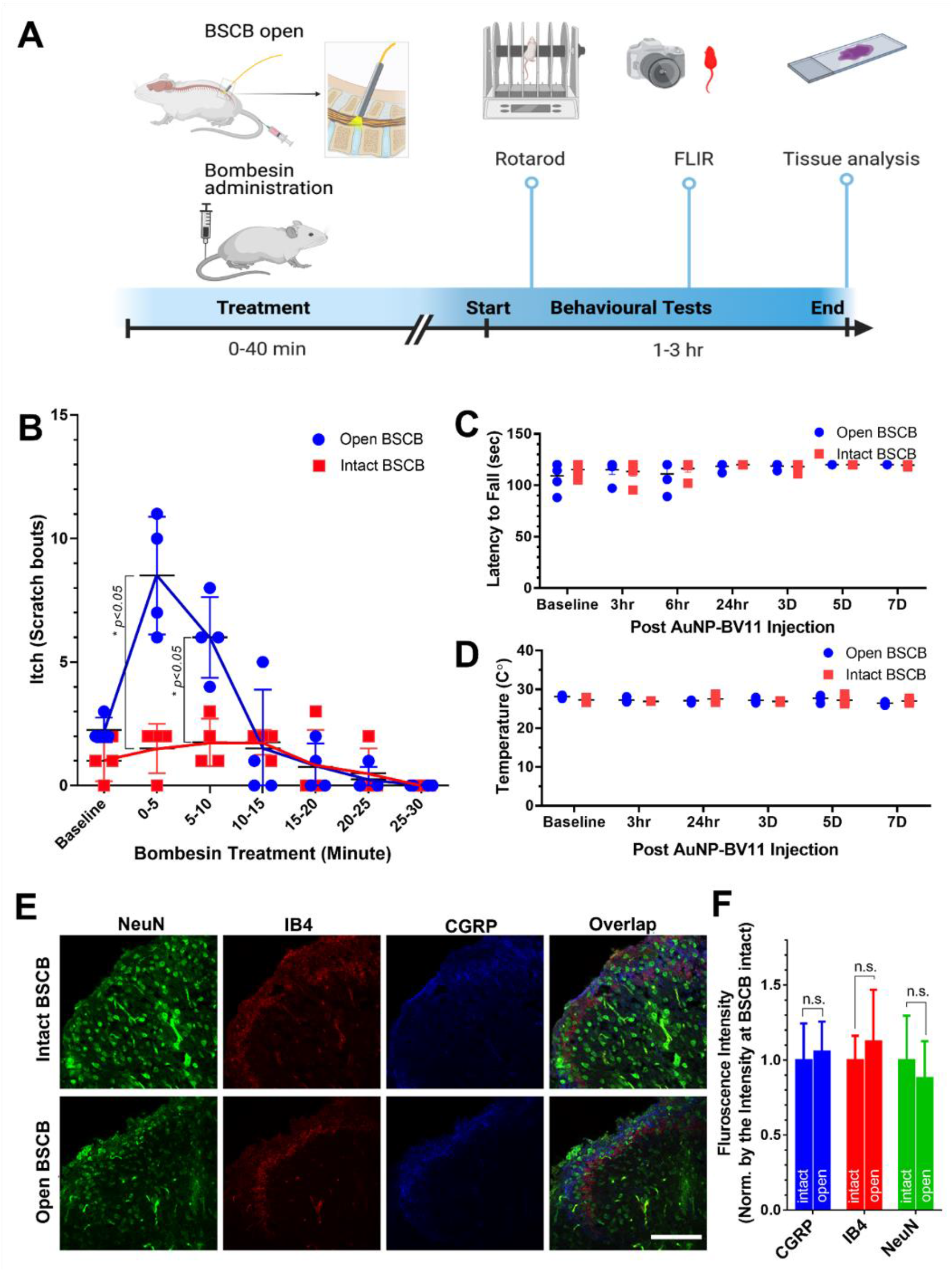
Local peptide delivery into the spinal cord and behavior modulation by Opto-BSCB. (A) Schematic and experimental workflow of Opto-BSCB modulation for peptide delivery. (B) Increased itching behavior in mice with open BSCB compared to control mice with intact BSCB. *N*=4 per group. (C-D) Impact of BSCB modulation on motor coordination and hind paw temperature. *N*=5 for each group. Data are expressed as mean ± SEM. Two-way ANOVA with Bonferroni correction for multiple comparisons. *P<.05, **P<.01. (E) IHC staining and (F) quantification of tissue from spinal cord regions with intact BSCB versus tissue from areas with open BSCB by NeuN (green), IB4 (red), CGRP (blue); Laser pulse number of 10 (5 Hz) at an intensity of 35 mJ/cm^2^ measured at the fiber tip. Scale bar: 100 µm.

We further investigated the impact of BSCB opening on specific cell type morphologies and numbers by IHC staining. We assessed major cell types in the spinal cord, including neurons (NeuN), astrocytes (GFAP), microglial cells (Iba1), IB4-positive sensory neurons (IB4), and calcitonin gene-related peptide positive sensory neuron (CGRP) (Fig. 3E; and Fig. S6). We observed no significant changes in NeuN, IB4, CGRP, or GFAP staining at the site with laser light compared to regions with no laser treatment (Fig. 3F; and Fig. S6). This result confirms that the BSCB opening and peptide delivery do not disrupt the neuronal integrity or cause astrogliosis or microglial reactivity changes, consistent with the previous section in dye permeability investigations.

## Discussion

One critical challenge in developing medicine for diseases/disorders in the spinal cord is the limited drug penetration due to the BSCB. Researchers have attempted to use focused ultrasound to open the BSCB and tested it in small and large animal models (6, 21). However, ultrasound waves will need to travel through and penetrate the solid bone and soft tissue to reach the spinal cord. There are challenges in bone attenuation of high-frequency ultrasound waves (22), thermal deposition (14), ultrasound focusing, and standing waves due to the presence of the vertebra and its unique bone structure (22) (9). Here, we report a method for the minimally invasive opening of the BSCB by an optical fiber. This method bypasses the vertebrae to directly deliver laser light to the surface of the spinal cord and has a potential path for clinical translation. The space between the two vertebrae and above the spinal cord is between 1 and 5 mm for humans (30), allowing access to the spinal cord with a large catheter up to 2.5 mm (23). In this context, it is possible to use a more significant optic fiber to deliver light to the spinal cord in humans. Indeed, optic fiber-based probes are under clinical trials to monitor spinal cord hemodynamics in large animals, such as pigs (24). Clinicians have also adapted spinal laser interstitial thermal therapy (LITT) for treating tumors in the spinal cord in human patients (25, 26). Compared to FUS, the Opto-BSCB method provides an effective and safe tool for delivering a wide range of molecules locally.

In terms of clinical translation to humans, researchers have used the intrathecal injection of drugs and implantation of pumps for spinal disorders and pain management (15, 27). Intrathecal injection (IT) involves the direct injection of therapies into the cerebrospinal fluid (CSF). IT is an attractive method for drug delivery due to the extended access of CSF to the CNS. However, limited IT-administrated therapies have been translated to the clinic, in part, due to limited distribution and rapid clearance. The fate of large-sized treatments (e.g., nanoparticles) after IT still needs to be investigated (28). The Opto-BSCB method takes advantage of the minimally invasive fiber-optic light delivery into the spinal cord and allows the penetration of molecules into the spinal cord parenchyma.

Opto-BSCB is most suitable for improving medical treatment of diseases affecting a local spinal region (e.g., spinal cord injury, tumor, pain). Many diseases (neurodegenerative diseases) can affect an extended length of the spinal cord, requiring opening the BSCB in a large volume of tissue. The current form of the fiberoptic device used in this study for delivering light is suitable for feasibly testing in small animals. Optic fibers can be integrated with mechanical platforms to send the laser light to multiple locations with a single injection through a side-view illumination (24). For improved precision, Opto-BSCB can be applied together with magnetic resonance imaging (MRI) and micro-CT to perform image-guided delivery. Optical fibers can also be integrated with advanced spinal endoscopes (white light and fluorescence) to complete molecular image-guided BSCB opening (29, 30). Furthermore, Opto-BSCB also has the potential to be used for studying neuromodulation in combination with chemogenetics (31) or optogenetics (16).

Since both gold nanoparticles and fiberoptic devices are in clinical trials (32, 33), this approach is valuable for clinical translation to improve therapeutic delivery to treat various diseases/disorders in the spinal cord. Many diseases/disorders affecting the spinal cord have genetic origins and are ideal candidates for gene therapy or protein replacement-based therapy. Significant progress has been achieved in the field of RNA and gene therapy. Notably, the messager RNA (mRNA) platform has been successfully used for developing advanced vaccines and has shown potential in treating other diseases (34). CRISPR gene-editing technology corrects single-point mutations with base editors (35) and large DNA sequence deletions, replacements, integrations, and inversions with twin prime editing (36). To carry these agents, viral and non-viral platforms are required to to facilitate them into targeted cells. Major gene-therapy carriers, including AAVs (< 20 nm) (37, 38) and lipid nanoparticles (LNP > 50 nm) (39, 40), are too large to penetrate the BSCB within the optimal therapeutic window. The Opto-BSCB approach may allow the delivery of AAVs and LNPs into the spinal cord.

One main limitation of the Opto-BSCB is the low light penetration due to strong light scattering in the visible wavelength range (532 nm). This issue is due to the myelinated axons enriching tissue geometry and organization in the spinal cord (41). We can overcome this limitation by shifting the optical window to the near-infrared (NIR) wavelength range (42). Nanoparticles with optical absorption in the NIR, such as gold nanorods, nanourchins, and nanostars, can be helpful for BSCB permeability modulation in deep tissue (43). Furthermore, the tiny diameter of the optic fiber used in this study also limits the penetration depth. At the same time, a larger beam size opens the BSCB in a large tissue region and depth (Fig 1). Therefore, we envision increasing penetration by using a large-diameter fiber (up to several mm) and side illumination to cover a greater depth in a large animal and human spinal cord (44, 45).

## Conclusion

We presented a novel optical approach for modulating BSCB permeability in the spinal cord and its ability to facilitate peptide delivery. This approach provides a short 24-hour window for drug delivery without significant side effects at the cellular and behavioral levels. This Opto-BSCB approach is compatible with fiberoptic light delivery for accessing the surface of the spinal cord with minimal invasiveness. We demonstrated a proof-of-the-concept for biological delivery by successfully delivering an itch behavior-causing peptide. Opto-BSCB has the potential for basic research and clinical utilities to improve drug delivery efficacy in the spinal cord.

## Materials and Methods

### Nanoparticle formulation and characterization

#### Materials and reagents

Gold (III) chloride, fluorescein isothiocyanate-labeled dextran (FITC-dextran, 70 kDa), Evans blue (EB), 4′,6-diamidino-2-phenylindole (DAPI), EZ-link biotin, hydroquinone, sodium citrate tribasic, Tween 20, Triton-X 100, sucrose, lectin-Cy5, Cy3-labelled streptavidin and Hoechst were purchased from Sigma-Aldrich. OPSS-PEG-SVA (3400 Da) and mPEG-thiol (1000 Da) were purchased from Laysan Bio, Inc. Li silver enhancement kit, donkey serum, goat serum, DyLight 594-labeled tomato lectin, gold reference standard solution, phosphate-buffered saline (PBS), borate buffer, 4% PFA in PBS 1x, nitric acid, hydrochloric acid, syringes, needles, and 20 kDa MWCO dialysis membrane were purchased from Thermo Fisher Scientific. All chemicals were analytical grade. Anti-JAM-A antibodies BV16 and BV11 were provided by Drs Elisabeth Dejana and Monica Giannotta at FIRC Institute of Molecular Oncology Foundation.

#### Gold nanoparticle (AuNP) formulation and conjugation

AuNPs were synthesized following a previously reported method and conjugated with antibodies. The size of AuNPs is controlled to around 48 nm in diameter and confirmed by transmission electron microscopy (TEM) and dynamic light scattering (DLS). For conjugation, BV11 or BV16, anti-JAM-A antibodies for our in vivo and in vitro models (respectively) were first diluted to 0.5 mg/mL in PBS 1x and transferred into 2 mM borate buffer (pH 8.5) with the final concentration of 0.05 mg/mL. Orthopyridyldisulfide-polyethylene glycol-succinimidyl valerate, average MW 3400 (OPSS-PEG-SVA), was dissolved in 2 mM borate buffer (pH 8.5) and quickly added to the diluted antibody solution at a 125:1 molar ratio. The mixture was mixed by vortex and kept on ice and gentle shaking for 3 hours. The resulting mixture was loaded into a 20 kDa MWCO membrane, placed into a 5 L tank of 2 mM borate buffer (stirring with a magnetic bar), and dialyzed at 4 °C overnight to remove free OPSS-PEG-SVA. The thiolated antibodies were mixed with concentrated AuNPs at a 200:1 antibody-to-NP ratio and reacted for 1 hour on ice. Next, mPEG-thiol was added to the reaction at a surface density of 6 PEG/nm^2^ for backfilling the empty surface space of AuNPs for 1 hour on ice to stabilize nanoparticles in the physiological environment. Finally, the functionalized nanoparticles were washed 3 times in 2 mM borate buffer and characterized by dynamic light scattering (DLS) and UV-vis spectroscopy. AuNP-PEG and AuNP-antibody were stored in 2 mM borate buffer (pH 8.5) at 4°C for use within 2 weeks to maintain the targeting activity.

#### Transmission electron microscopy (TEM) of AuNPs

Three drops (2.5 μL per drop) of AuNP nanoparticle suspension (200 μg·mL^−1^) were deposited on a formvar/carbon film-coated copper grid (Ted Pella, Inc.) and dried in a desiccator for at least 2 hr or overnight at room temperature. Afterward, TEM images of nanoparticles were captured on a Field Electron and Ion Company (FEI) Tecnai Transmission Electron Microscope. TEM imaging was carried out by TEM-JEOL 1400+ (JEOL).

#### Dynamic light scattering (DLS) of AuNPs

An aliquot of the abovementioned AuNP suspension was diluted in 2 mM borate buffer to a final concentration of 200 μg·mL−1. Then, DLS measured the hydrodynamic diameters of diluted nanoparticles in solutions on a Malvern Nano-ZS Particle Sizer (Malvern Panalytical Ltd.).

#### UV-Vis-NIR spectroscopy of AuNPs

The UV-Vis-NIR absorption spectrum of the nanoparticle suspension was measured by a Cary 6000i spectrophotometer (Agilent) with a total path length of 1 mm, background-corrected for contribution from water and the cuvette. The measured range was 200 to 800 nm.

### Administration of nanoparticles and bombesin

#### Administration of nanoparticles

AuNP-PEG or AuNP-BV11 were centrifuged at 1300 G for 30 minutes at 4°C and then dispersed into 2 mM borate buffer at the desired concentration. For injection into animals, 2 mM borate buffer is removed by centrifugation and redispersed in saline. 90 μL of nanoparticle suspension was intravenously administrated at the dose of 18.5 mg/kg nanoparticles for both AuNP-PEG and AuNP-BV11.

#### Administration of bombesin

A stock solution of [Lys^3^]-Bombesin (Sigma Aldrich, Cat# B1647) was prepared in sterile saline (2 mg/mL) on the day of the experiment and serially diluted to 50 μg/mL, then intravenously infused into the tail vein with a polyethylene tubing (PE-20) catheter, all catheter lines were flushed using 50 μL of sterile saline or PBS1x buffer by a Gastight 1700 Series Syringe (Hamilton Cat# 80901). The total injection volume is limited below 250 μL for an individual mouse with a bodyweight < 25 g.

### Intrathecal device implantation and BSCB modulation

#### Intrathecal injection

As previously described, a 27.5-gauge butterfly needle was injected into the intrathecal space (46). A 50-μm core optic fiber was inserted through the inner channel of the butterfly cannula to reach the intrathecal space. The optic fiber is coupled to the picosecond (ps) laser to excite the AuNPs located in blood vessels. The optical fiber is pointed to the spinal location of interest after laminectomy.

#### Opto-BSCB modulation

The animal was anesthetized with 2-3% isoflurane and intravenously administrated with 18.5 mg/kg AuNP-BV11. The 532 nm picosecond (ps) laser (5 Hz, intensity 30-40 mJ/cm^2^) at an optimal pulse of 1-100 was applied to excite the AuNPs in blood vessels. Tracers, EZ-link biotin (2 mg/mL in saline, 100 μL) and FITC-dextran (40 mg/mL in saline, 100 μL), Cy5-lectin (2% in PBS, 100 μL), or Evans blue (2% in PBS, 100 μL) were injected into the tail vein at different time points post-laser stimulation. After 1 hour, mice were transcardially perfused with 25 mL of cold PBS 1x to remove the free-floating intravascular dye within the blood, followed by 25 mL 4% PFA in PBS 1x. The spinal cord and other organs were extracted for staining and imaging. Particularly, the spinal cord was post-fixed in 4% PFA in PBS 1x overnight, then dehydrated in 30% sucrose in water for another night for IHC staining. To quantify the distribution of tracers, we cut fixed tissue sections with 30-μm thickness at −20°C on a cryostat. Sections were mounted on glass slides and fluorescently imaged by a slide scanner (VS120, Olympus). Sections collected from mice injected with EZ-link biotin, FITC-dextran, and Cy5-lectin, were stained by Cy3-labelled streptavidin (1:200) to detect EZ-link biotin and Hoechst to label nuclei and imaged.

### Animals, biodistribution, tissue staining, and analysis

#### Animal

Adult male C57BL/6 mice (8 weeks old, 22-25 g) were ordered from Charles River Laboratories. Animal protocols were approved by the Institutional Animal Care Use Committee (IACUC) of the University of Texas at Dallas.

#### Biodistribution

To confirm whether AuNPs can be distributed in the spinal cord and examine the clearance time in mice, we perfused the animal and collected major organs at 1 hour and 2 weeks after IV injection of AuNPs. The tissue was then digested in aqua regia (1 part nitric acid, 3 parts hydrochloric acid) for 3-7 days and centrifuged at 16000 G for 10 minutes. The supernatant was collected and diluted approximately 40x in water or to fit within the range of standard curve concentrations for measurement. The gold concentration was analyzed by inductively coupled plasma mass spectrometry (ICP-MS). To microscopically examine AuNP-BV11 distribution, major organs were excised for histology staining using silver enhancement reagents. We performed paraffin processing of tissues 20 µm thick, followed by deparaffinization steps. The sections were then rinsed with Milli-Q water and incubated with Li silver enhancement developer for 20 minutes at room temperature. After washing in water, the treated tissues were counterstained with hematoxylin for 5 minutes, followed by dehydration through 100% ethanol and clearance by xylenes. Finally, the tissues were covered with synthetic mounting media and imaged under bright field microscopy. ICP-MS: Mouse organs were weighed, and aqua regia was added to each organ to a ratio of 5 mL aqua regia per g of organ tissue. Organs were digested for >3 days, then the digested liquid was centrifuged at the maximum speed for 10 minutes, the supernatant was collected, and the supernatant was diluted further to a concentration of 2-5% acid. Then, organs predicted to have large concentrations of particles (spleen, liver, lung) were diluted 10 times. These diluted acid solutions were then used for ICP-MS readings.

#### Immunocytochemistry

To detect if functionalized AuNPs can target JAM-A, we performed ICC staining with an in vitro BBB model, hCMEC/D3 cell (human cerebral microvascular endothelial cell) monolayer. AuNPs were conjugated with BV16 (anti-JAM-A antibody) for targeting. D3 cells were seeded at a density of 30000 cells/cm^2^ and cultivated for one week to form a monolayer at 37 °C and 5% CO2. Cellular monolayers were then incubated with 0.5 nM AuNP-BV16 for 30 minutes, followed by fixation for 5 minutes in pure methanol. The samples were then incubated with a blocking buffer (5% donkey serum in PBS 0.05% Tween), and a secondary antibody (A21202, ThermoFisher) incubated at room temperature (RT) for 1 hour at each step. Finally, the monolayers were stained with Hoechst dye for 7 minutes in the dark to label the nucleus. The samples were washed in PBS x1, three times between each step. FV3000RS confocal microscopy was used to take images.

#### Immunohistochemistry (IHC)

Mouse spinal cord was snap-frozen on dry ice and stored at −20 °C before processing into 40-μm thick slices using a cryostat. To stain Iba1, GFAP, Ib4, and NeuN, the mouse tissues were dehydrated in a 30% sucrose solution overnight. The dehydrated tissues were sliced to 40-μm thickness on a freezing cryostat into 0.03% sodium azide in PBS 1x solution. Sections were washed in PBS 1x three times to remove the cryoprotectant solution completely. After blocking (20% normal goat or donkey serum with 0.1% Triton X-100 in PBS) for 1 hour at room temperature, free-floating sections were incubated with primary antibodies: anti-Iba1 (AB48004, Abcam), anti-GFAP (RB087A0, Fisher Scientific), anti-IB4 (I21412, Invitrogen), for two nights at 4°C. Secondary antibodies were then added, followed by incubation with DAPI solution. For staining of NeuN, the mice were perfused with PBS and 4% PFA at 5 mL/min, followed by post-fixing in 4% PFA overnight. We sliced the tissue to a thickness of 40 μm on a cryostat at −20°C after dehydration in a 30% sucrose solution overnight. The sections were washed in milli-Q water at 37°C for 5 minutes and incubated with pepsin solution for antigen retrieval. The treated cells were then incubated with blocking buffer at room temperature for 1 hour and primary antibodies (anti-NeuN (NC1284461, Fisher Scientific)) overnight at 4°C. Secondary antibodies were applied for 1 hour at room temperature, followed by incubation with a 1:2000 volume ration of Hoechst in PBS solution.

#### Imaging and analysis

Imaging was performed with a slide scanner (VS120, Olympus). Fluorescent intensity was analyzed by CellFluo software for each section. For each mouse, total fluorescence intensity was summed from individual sections.

### Behavior studies and analysis

#### Forward-Looking Infrared (FLIR) System

The temperature of the animals’ hind paws was used as a measure of potential paw inflammation. All testing was performed in a climate-controlled room with an ambient temperature of 21 ± 2°C. Animals were acclimated in the testing room for 1 hour before testing. Colorized Infrared thermograms that captured the non-glabrous surface of the animal’s hind paws were obtained using a FLIR T-Series Thermal Imaging Camera (T650SC). The thermograms were recorded before experimental treatment, at 3, 6 hours, 24 hours, day 3, and 7 days after nanoparticle delivery. Thermogram analysis was performed using the Windows-based PC application of the FLIR system. For each thermogram image, an ROI was drawn on the plantar surface of both hind paws, and the mean temperature was recorded from the average of each pixel by a blinded analyst to the conditions.

#### Rota Rod

For an assessment of motor coordination after opening the BSCB, we measured animals on the Rota Rod (IITC Life Sciences), accelerating from a starting speed of 10 rpm to 30 rpm in 60 seconds (s). Animals were given a rest time of 180 s between each trial. The end of a trial was noted when a mouse fell off the rod or when it reached the 120-second time point. A total of 3 trials were recorded per animal, and the average time was recorded.

#### Itch Assay

One day before bombesin (BBS) administration, mice were habituated in acrylic boxes for 1 hour. Cameras were placed in front and behind the mice, with recordings taken simultaneously for 30 minutes. The following day, mice received the AuNP-BV11 IV injection and activation with an IT administered 532 nm laser, then injected IV with 50 µg/mL of BBS following BSCB modulation and allowed to habituate for 30 minutes before recording itch behavior. Cameras were placed in front and behind the mice and simultaneously recorded for 30 minutes. Itch bouts were measured as previously described by (46). Scores were measured by an experimenter blinded to the conditions.

### Statistical Analysis

All the data were plotted and analyzed with Origin 2020. For the in vivo biodistribution study using ICP-MS, one-way ANOVA was used to compare the accumulation difference in main organs between targeting and non-targeting gold nanoparticles. For the kinetics study using different tracers, a Student’s t-test was used to compare the total fluorescent intensity between different time points. We performed a Student’s t-test to analyze confocal images of tissue and compare the difference between laser and no laser treatment.

## Acknowledgments

We are grateful to grant support funded by the Department of Defense (DOD: W81XWH2110219, to Z. G), National Institutes of Health (NIH: NS065926, to T. P.), American Hospital Association (AHA: 19CSLOI34770004, to ZQ), and Cancer Prevention and Research Institute of Texas (CPRIT: RP190278, to ZQ).

## Author’s contribution

ZG, ED, TP, and ZQ conceived the concept and designed the experiments; ZG, ED, TL, XQ, QC, JW, and SK performed experiments; MG, ED, prepared and provided BV11 and BV16 antibodies. RB provided training on animal surgery and insight into the study design; ZG and ED analyzed data; ZG, TP, ZQ. supervised the study. All authors reviewed the data and wrote the manuscript.

---

**Figure S1.**
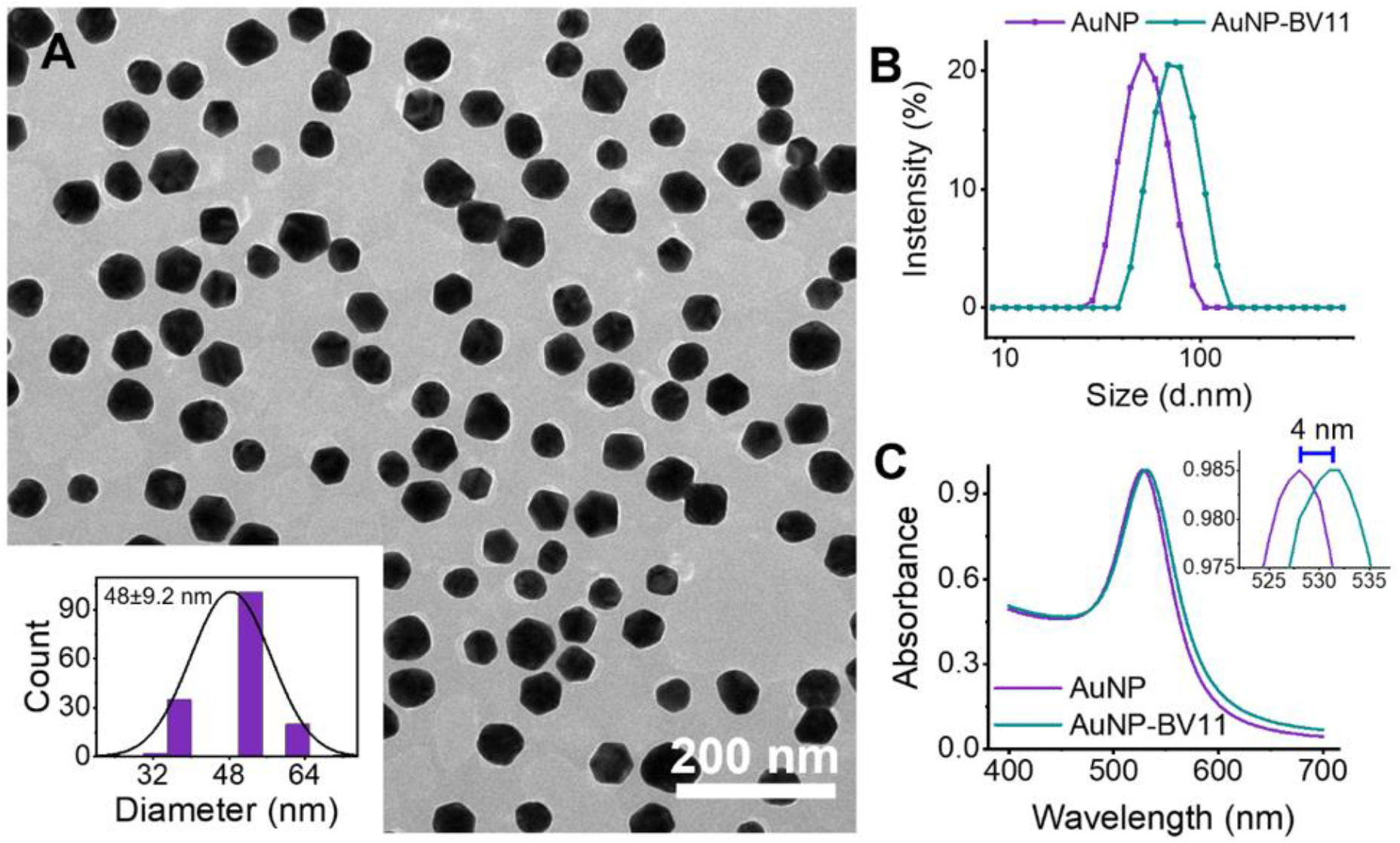
Characterization of JAM-A targeted AuNPs. (A) TEM of AuNP-BV11. Scale bar: 200 nm. Insert shows the diameter distribution of AuNP-BV11 based on the TEM image. (B) The hydrodynamic size of AuNP and AuNP-BV11. (C) The absorbance of AuNP and AuNP-BV11.

**Figure S2.**
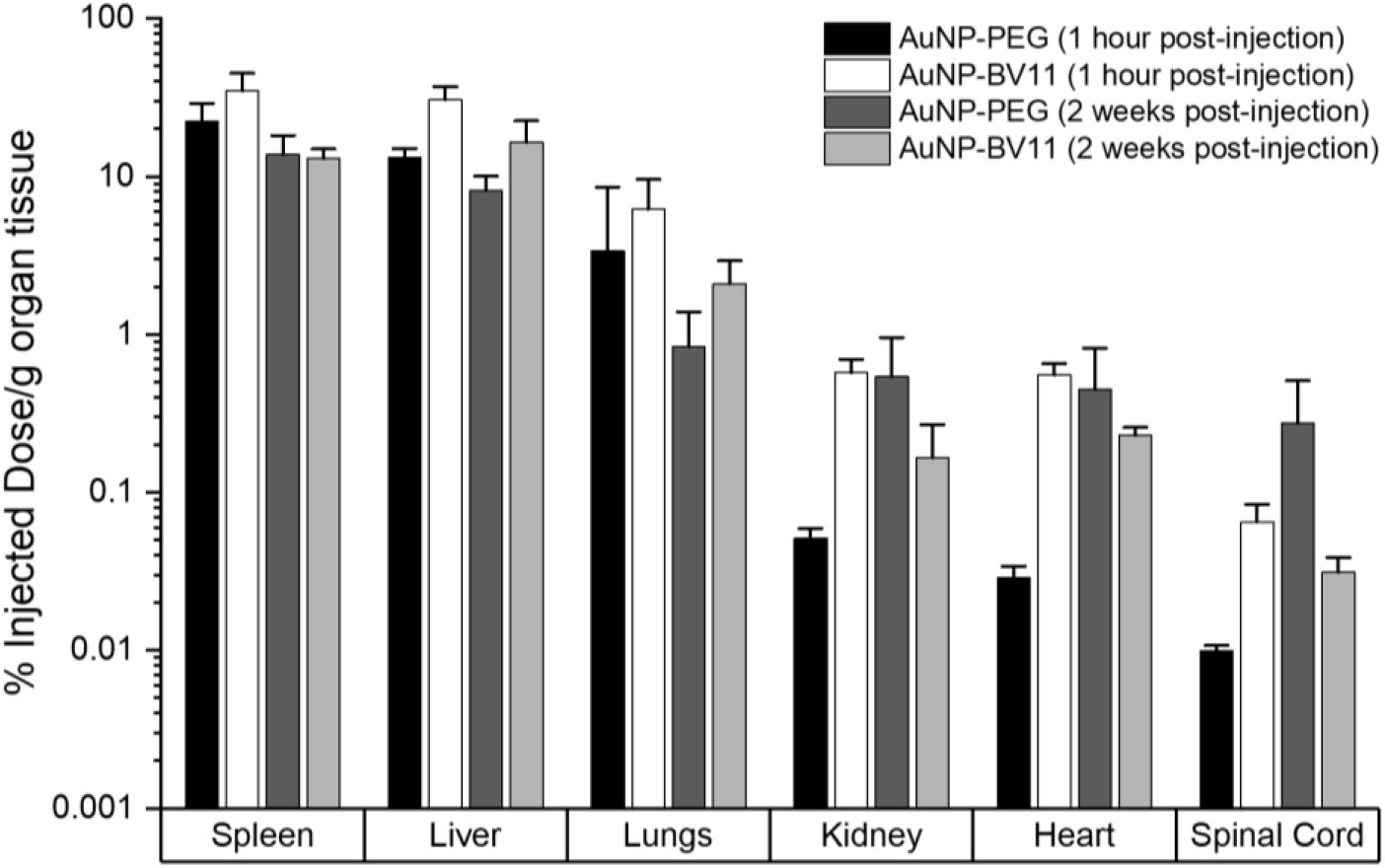
Biodistribution in major organs. Biodistribution of AuNP-BV11 and AuNP-PEG at 1 hour and 2 weeks post-injection. The animal was intravenously administrated 18.5 mg/kg AuNP-BV11 or AuNP-PEG. *N*=3 for each group.

**Figure S3.**
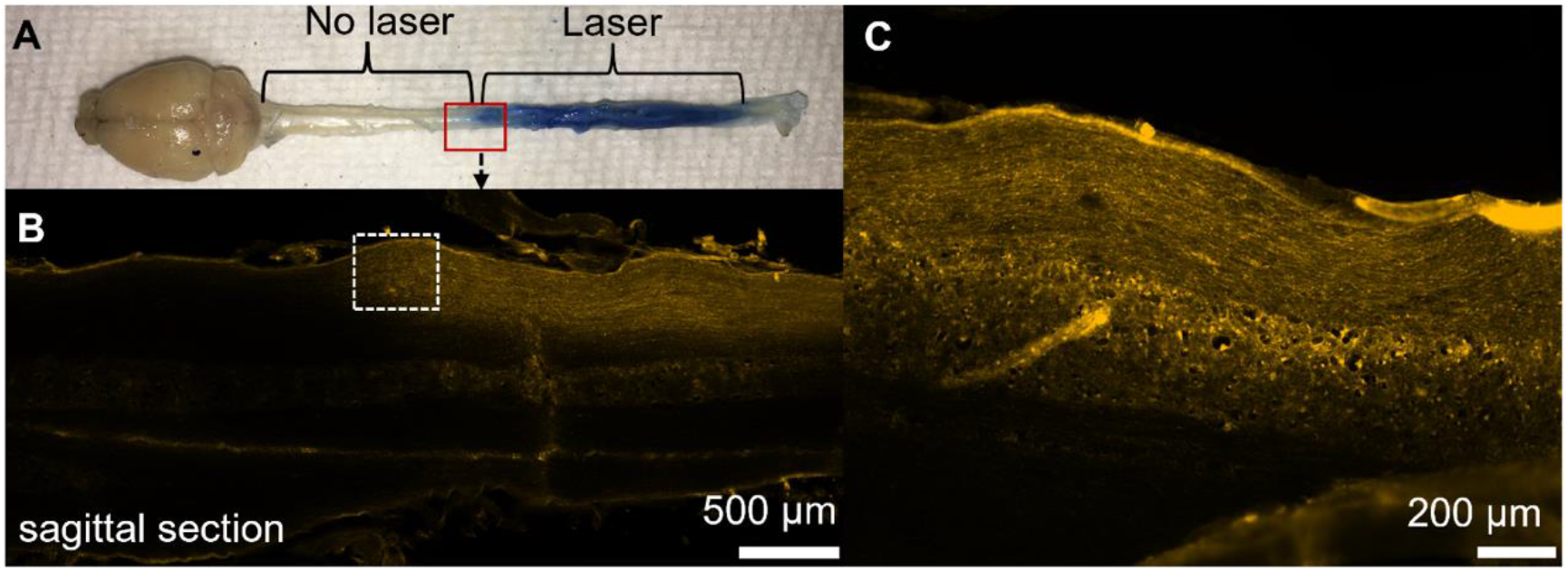
BSCB modulation in mice with a laser beam penetrated through the intact skin/bone tissue. (A) The blue-colored area indicates EB extravasation in the spinal cord region, suggesting BSCB opening by the laser light stimulation. In contrast, no EB was detected in areas that had no laser light stimulation. (B) Fluorescent image of the sagittal section of the spinal cord tissue with the left half without laser irradiation and the right half with laser irradiation. Scale bar: 500 µm. (C) Zoom-in view of the dashed square area in B. Scale bar: 200 µm. Laser conditions: 40 mJ/cm^2^ and 100 pulses scanned through the intact tissue. The laser beam size is 6 mm in diameter.

**Figure S4.**
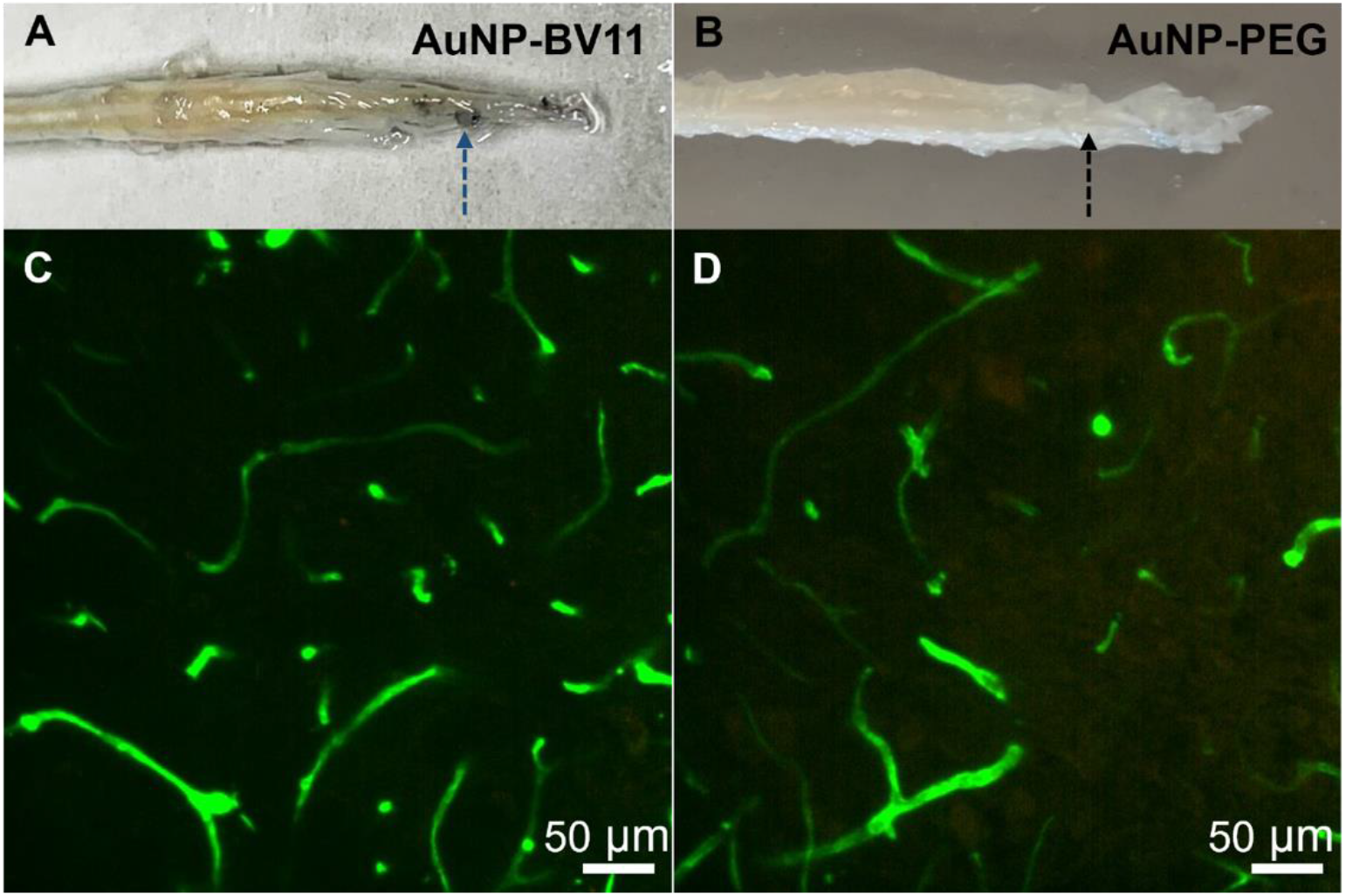
Comparison of BSCB modulation using AuNP-BV11 and AuNP-PEG. (A) EB extravasation into the spinal cord in the mouse injected with AuNP-BV11. The blue spot area suggests BSCB opening. (B) No EB extravasation was observed in the mouse injected with AuNP-PEG. (C-D) High-resolution fluorescent images show no significant difference in the blood vessel morphology between tissue (C) with BSCB opening versus (D) without the BSCB opening. Blood vessels are labeled by IV injection of lectin. Animals were sacrificed 30 minutes after laser stimulation. Scale bar: 50 µm.

**Figure S5.**
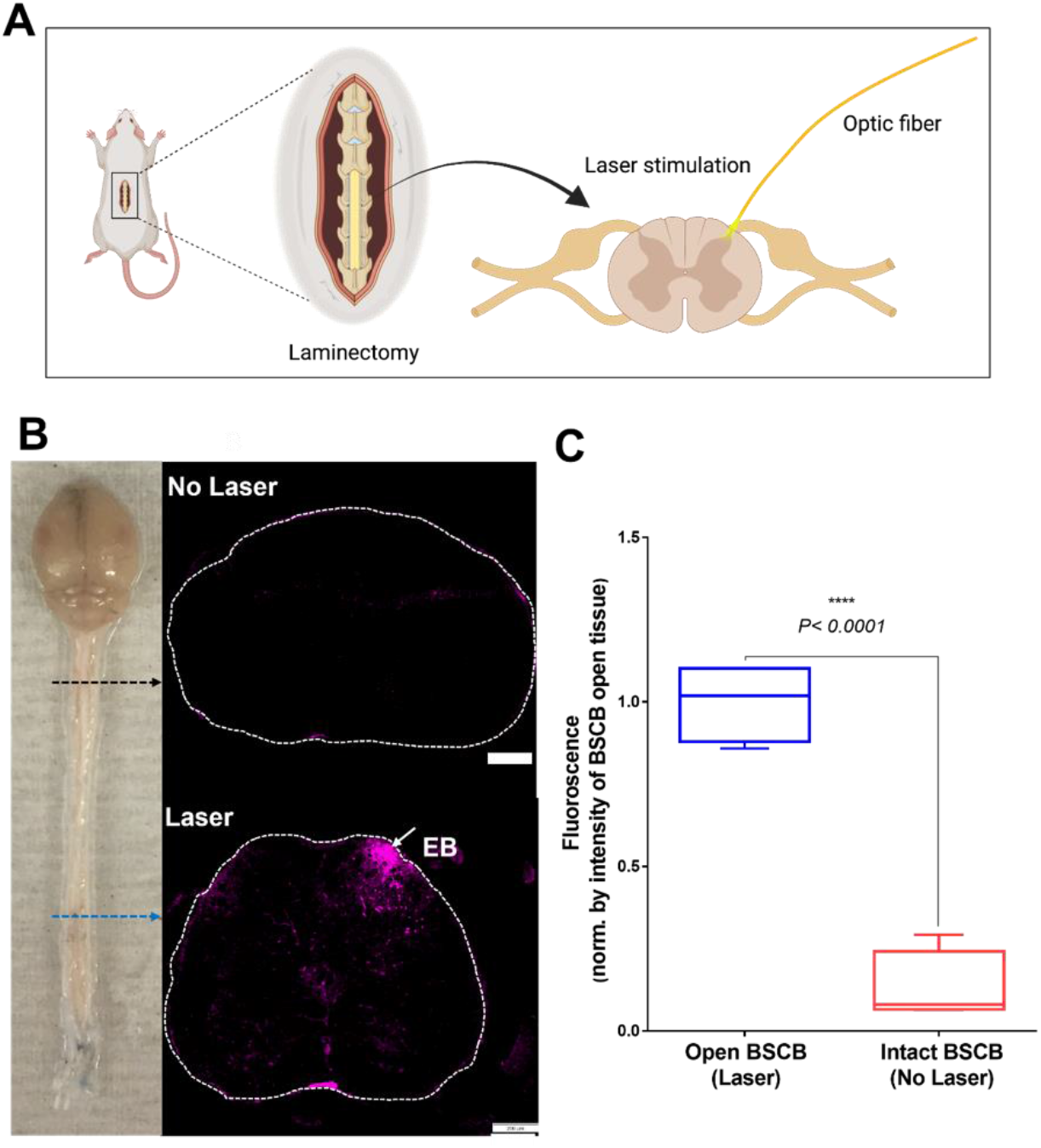
Local Opto-BSCB modulation with optic fiber (200 µm dinamter). (A) Schematic representation of the BSCB modulation with optical fiber. (B) The image of a mouse CNS shows the EB dye penetrated spinal tissue regions that received laser stimulation. A laser pulse number of 5 (5 Hz) at an intensity of 35 mJ/cm^2^ measured at the fiber tip was applied to the lumbar region. Fluorescence imaging of the cross-section of spinal cord tissue collected from areas with and without laser stimulation. Scale bar: 200 µm. (C) Quantitative comparison of the fluorescence in the spinal cord with BSCB open side vs. intact side, 7.7-fold higher fluorescence in the open BSCB tissue side as compared to intact BSCB tissue side (fluorescence is indicated as mean ± SD, normalized by the mean intensity at BSCB open side, *N*=4, p<0.0001, T-test).

**Figure S6.**
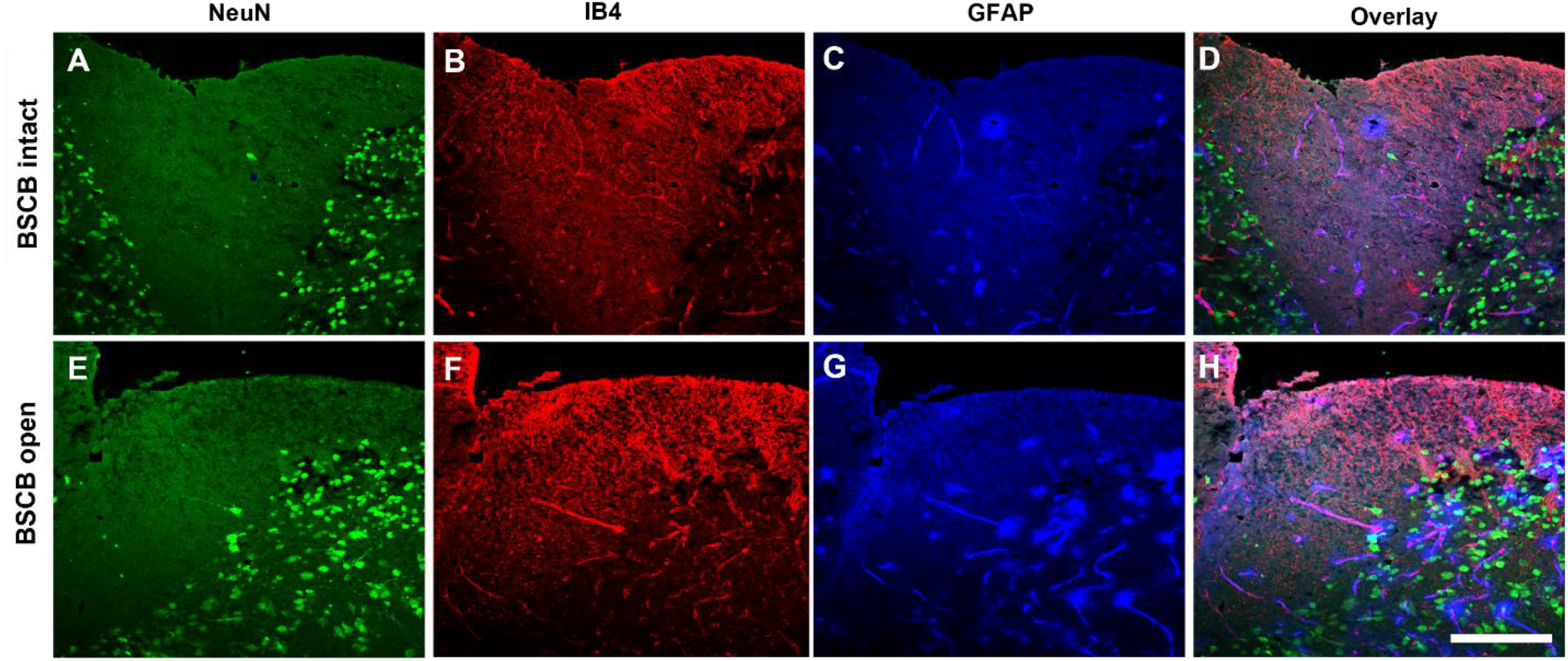
Impact of BBS delivery on the major cellular morphology. IHC staining of spinal cord tissue after bombesin (BBS) delivery in mice with (A-D) BSCB intact versus (E-H) BSCB open. A and E: NeuN; B and F: IB4; C and G: GFAP; and D and H: overlay of A-C, E-G, respectively. Scale bar: 100 µm.

(Uploaded separately as “Supplementary Video S1”)

**Video S1**. An example of spinal itch bouts induced after BBS delivery in Open BSCB vs. Intact BSCB animals. From left to right in the front row: boxes 1 and 2 have BSCB intact, while boxes 3 and 4 have BSCB open

